# NMR as a Video-(game): Constructing Super-Resolution Cross-peak Trajectories in Protein Spectroscopy

**DOI:** 10.64898/2026.03.19.712888

**Authors:** Chng Jing Hang, Yevheniia Kuznietsova, Mikhail Fillipov, Konstantin Pervushin

## Abstract

High-resolution multidimensional NMR spectroscopy of proteins remains limited by long acquisition times, sensitivity constraints, and severe peak overlap, particularly for larger systems. Conventional 3D and higher-dimensional experiments trade experimental efficiency for resolution, while post-acquisition analysis often becomes the dominant bottleneck. Here, we present a new framework that redefines both how NMR experiments are constructed and how they are executed and analyzed, by treating an AI agent-controllable series of 2D spectra as a spatiotemporal dataset analogous to a video. Our approach is based on temperature-dependent series of reduced-dimensionality 2D HSQC and novel RDL-TROSY experiments, in which each 2D [^1^H,^15^N] cross-peak is controllably shifted and split in proportion to the ^13^C chemical shift of the J-coupled carbons. We propose treating a variable-temperature (VT) series as a pseudo-temporal video sequence in which each cross-peak traces a physically motivated trajectory through frequency space. The proportionality coefficient (α) of this reduced-dimensionality encoding is systematically and programmatically varied together with the temperature providing full control for constructing optimal cross-peak trajectories. As a result, individual resonances follow predictable, spectral acquisition time-controllable trajectories in the 2D spectral plane across the series, which can be executed by an autonomous AI agent directly interacting with the NMR GUI layer. Each spectrum represents a single “frame,” while temperature and RD controls serves as the temporal dimension. We describe two complementary super-resolution strategies: a cross-peak model-independent approach based on the deep-learning video super-resolution that leverages temporal redundancy to sharpen per-frame peak shapes, and a model-based approach that derives the exact mathematical form of the peak trajectories and uses it to design acquisition schedules that render individual peak paths maximally distinct and amenable for algorithmic deconvolution. As a result, we obtained full backbone resonance assignment in the wide temperature range (279–315 K) with one degree Kelvin resolution in a test protein in an automatic manner in the time frame typically required for collection of a single 3D NMR dataset.

## Introduction

Backbone resonance assignment is the prerequisite for all quantitative protein nuclear magnetic resonance (NMR) spectroscopy (1, 2), and its primary obstacle is spectral overlap. Conventional solutions add indirect dimensions, spreading peaks across additional frequency axes that form orthogonal spectral dimensions after Fourier transformation, at the cost of linearly increased acquisition time, which becomes prohibitive for experiments requiring large numbers of indirect time-domain sampling points (3). Variable-temperature (VT) NMR offers an independent additional axis. Backbone amide ^1^H, ^15^N, and ^13^C_α_ chemical shifts all respond to temperature in a residue-specific manner (3–5), reflecting local hydrogen bond geometry and backbone dynamics. The temperature coefficient ∂δ_HN_/∂T is linear over a wide temperature range (269–369 K) (5) and takes values spanning roughly −16 to +2 ppb K^−1^ for amide protons, with buried residues showing attenuated sensitivity (5). Since this linearity is well established, we acquire 40 spectra across a temperature range falling within this window, at 1 K intervals. This generates a pseudo-temporal sequence in which each temperature step corresponds to a time point, and each cross-peak traces a distinct, predictable path through frequency space, encoding the differential change in chemical shift drift between successive spectra. Two peaks overlapping at one temperature are often separated at another temperature or reduced-dimensionality (RD) settings; the question is how to exploit this information systematically across the full series of acquired spectra rather than discard it through frame-by-frame analysis.

The conceptual core of this paper is the bridging of VT-NMR to a computer vision framework, in which a non-spectroscopic variable (temperature) as well as signal acquisition time-controlled variables are exploited to resolve physics motivated spectroscopic observables such as chemical shifts or selected NOEs. The CV field has developed extensively, with deep learning-based methods now achieving robust detection and tracking of objects across successive frames, exemplified by YOLO (7) for detection and DeepSORT (8) for multi-object tracking. Each spectrum is a 2D frame; each point of the spectral intensity grid can be viewed as a pixel; the temperature and RD axis is a pseudo-temporal axis; and cross-peak drift is a motion field. This bridge has direct operational consequences: resolving transiently occluded peaks becomes a problem of propagating information across the temporal dimension, for which optical flow, recurrent feature propagation, and super-resolution are directly applicable.

Our approach differs from that of Shchukina et al. (9), who sample temperature and indirect evolution times jointly within a single acquisition, necessitating correction for signal non-stationarity arising from temperature drift during acquisition. In contrast, we acquire a fully stationary RD-HSQC or RDL-TROSY spectrum at each discrete temperature step; each frame is recorded independently and is therefore free from non-stationarity artefacts. The resulting challenge lies in linking and exploiting this discrete series of stationary spectra. Trainor, K et al. addressed a related problem by implementing peak tracking across temperatures to determine temperature coefficients, showing that the chemical shifts of ^1^H and ^15^N nuclei exhibit an approximately linear dependence on temperature. In our work, we extend this concept by exploiting the temperature-dependent chemical shift dynamics of three types of spins within a single spectral series, introducing a temperature-based encoding that captures pseudo-temporal correlations between spectra.

## Results

The experiment of choice is the reduced-dimensionality heteronuclear single quantum coherence (RD-HSQC) spectroscopy (10, 11). In addition, we introduced novel Reduced Dimensionality L-TROSY (RDL-TROSY) experiment maximizing achievable spectral resolution in all spectral dimensions. In this experiment, the ^13^C_α_ evolution time is linearly co-incremented with the ^15^N evolution time during the indirect dimension, so that the apparent ^15^N chemical shift is a linear combination of the true ^15^N and ^13^C_α_ offsets (10). In a conventional ^1^H– ^15^N HSQC, two residues with similar ^1^H and ^15^N shifts overlap regardless of their ^13^C_α_ differences. The RD-HSQC encodes the ^13^C_α_ frequency offset into the detected ^15^N dimension via a constant-time evolution element with a mixing coefficient β = SWH_N_ /SWH_C_, producing two symmetrically displaced cross-peaks per residue—the S^+^ and S^−^ components—at apparent ^15^N positions:

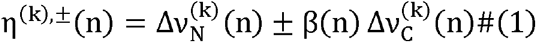

where β(n) = SWH_N_(n)/SWH_C_(n) is the frame-dependent mixing ratio (ratio of ^15^N to ^13^C_α_ spectral widths, both in Hz), k indexes residues, and n indexes temperature frames. The original two heteronuclear coordinates are recovered exactly by sum-and-difference inversion:

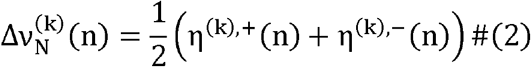

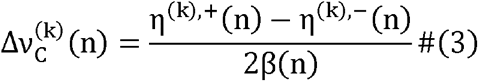

The encoding is lossless when β ≠ 0. By modulating SWH_C_(n) across the temperature series and thereby varying β(n), residues with different ^13^C_α_ offsets are displaced by different amounts at each frame, converting latent ^13^C diversity into a time-varying differential displacement that renders otherwise parallel trajectories divergent and individually trackable. Note that modulating SWH_N_(n) achieves the same effect; in practice SWH_C_(n) is the more convenient variable because the ^13^C_α_ window is narrower and more easily stepped without aliasing.

Two complementary resolution strategies follow from this reframing. The cross-peak model-independent strategy adopts video super-resolution (12, 13) to propagate spectral features bidirectionally across temperature steps without assumptions about the specific cross-peak point spread function (PSF). The cross-peak model-based strategy uses the exact master equation for RD-HSQC trajectories to design acquisition schedules that maximize trajectory separability. Together they operate at the two levels where the video reframing has traction: the reconstruction of per-frame peak profiles, and the design of the frame sequence itself.

### Master equation for RD-HSQC trajectories

Let the chemical shift offsets from carrier for residue k at temperature step n be 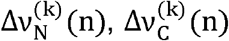 and 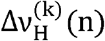 for ^15^N, ^13^C_α_, and ^1^H_N_ respectively. The temperature dependence of backbone amide shifts is approximately linear over moderate ranges (4):

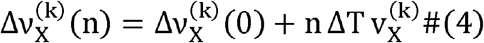

where ΔT is the temperature increment per step and 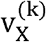 is the per-residue drift velocity (Hz K^-1^) for nucleus X. All three nuclei share a common carrier drift component that displaces the entire peak cloud coherently:

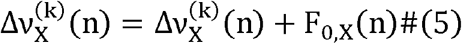

where F_0,X_ (n) is a shared, frame-dependent carrier correction (arising, for example, from deuterium lock drift or thermal expansion of the probe).

The complete observed peak positions in the 2D RD-HSQC spectrum are then

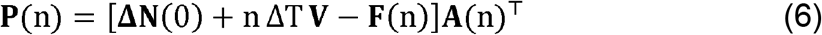

where **ΔN** (0) ∈ ℝ^Mx3^ is the matrix of initial offset coordinates for all M residues (columns ordered as Δ*v*_N,_ Δ*v*_C_, Δ*v*_H_), **V** ∈ ℝ ^Mx3^ is the per-residue drift velocity matrix, **F**(n) is a rank-1 broadcast matrix encoding shared carrier drift, and:

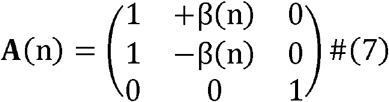

Each row-pair of **P** (n) gives the observed (η^+^,η ^-^,Δ*v*_H_) coordinates of one residue at step n. Because **F**(n) is rank-1 and maps identically to all residues, it cancels exactly in all pairwise peak differences and has no effect on inter-residue separability; its sole role is to keep the peak cloud centered within the spectral window. The drift velocity matrix **V**, drawing per-residue coefficients from distributions well-characterised for soluble proteins (4), is the fundamental source of trajectory distinctiveness. The RD projection matrix **A**(n) translates ^13^C_α_ offset diversity into differential separation in the mixed ^15^N dimension.

### Complementary S^+^ and S^−^ spectral planes as independent channels

The spin-state-selective editing of the RD-HSQC yields two spectral planes, S^+^ and S^−^, each containing one cross-peak per residue. The construction of planes is achieved by acquiring x and y quadrature components of the evolving ^13^C_α_ signal, *S*_in_ and *S*_anti_, then computing S^±^ = *S*_in_ ± *S*_anti_ (10, 11). The resulting S^+^ and S^−^ spectra function as two independent video sequences whose motion fields are coupled by Eq. (1). For a correct S^+^/S^−^ pair, the sum 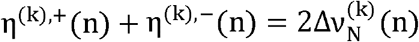 must vary smoothly with temperature, providing a stringent cross-validation constraint: spurious pairings produce erratic sums, correct pairings produce smooth ^15^N trajectories. Because S^+^ and S^−^ peaks are spatially separated by 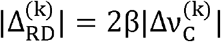, peaks occluded in one channel may be resolved in the other. The S^+^/S^−^ splitting magnitude is directly proportional to the ^13^C_α_ offset from the ^13^C carrier. Residues near the carrier (Δ*v*_C≈_0) produce nearly coincident S^+^/S^−^ pairs and therefore contribute little discriminating information from the RD dimension; the acquisition schedule should set the carrier to maximise the spread of 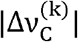 values across all residues.

Figure 1 shows overlay of 40 spectral planes of S^+^ and S^−^ RD-HSQC measured with 76 amino acid test protein (Ubiquitin). Placing the ^13^C_α_ carrier at 52 ppm (near the centre of the C_α_ chemical shift range for amino acids) achieves adequate coverage for most residues in the protein. The ^13^C spectral width SWH_C(n) and the coupled ^1^□N spectral width SWH_N(n) were modulated as linear functions of the frame index n across the 40-step temperature ramp, subject to the reciprocal-sum constraint 1/SWH_N(n) + 1/SWH_C(n) = C_const (Eq. 8), which ensures that β(n) evolves in a controlled, monotonic fashion. This frame-by-frame modulation of β(n) actively steers the trajectory of each S^+^/S^−^ peak pair in the mixed ^1^□N dimension: residues with larger |Δ*v*^(^□^)^_C| are displaced more strongly at each frame, converting latent ^13^Cα chemical shift diversity into a time-varying differential displacement that renders otherwise parallel trajectories divergent and individually trackable across the series.

**Figure 1.**
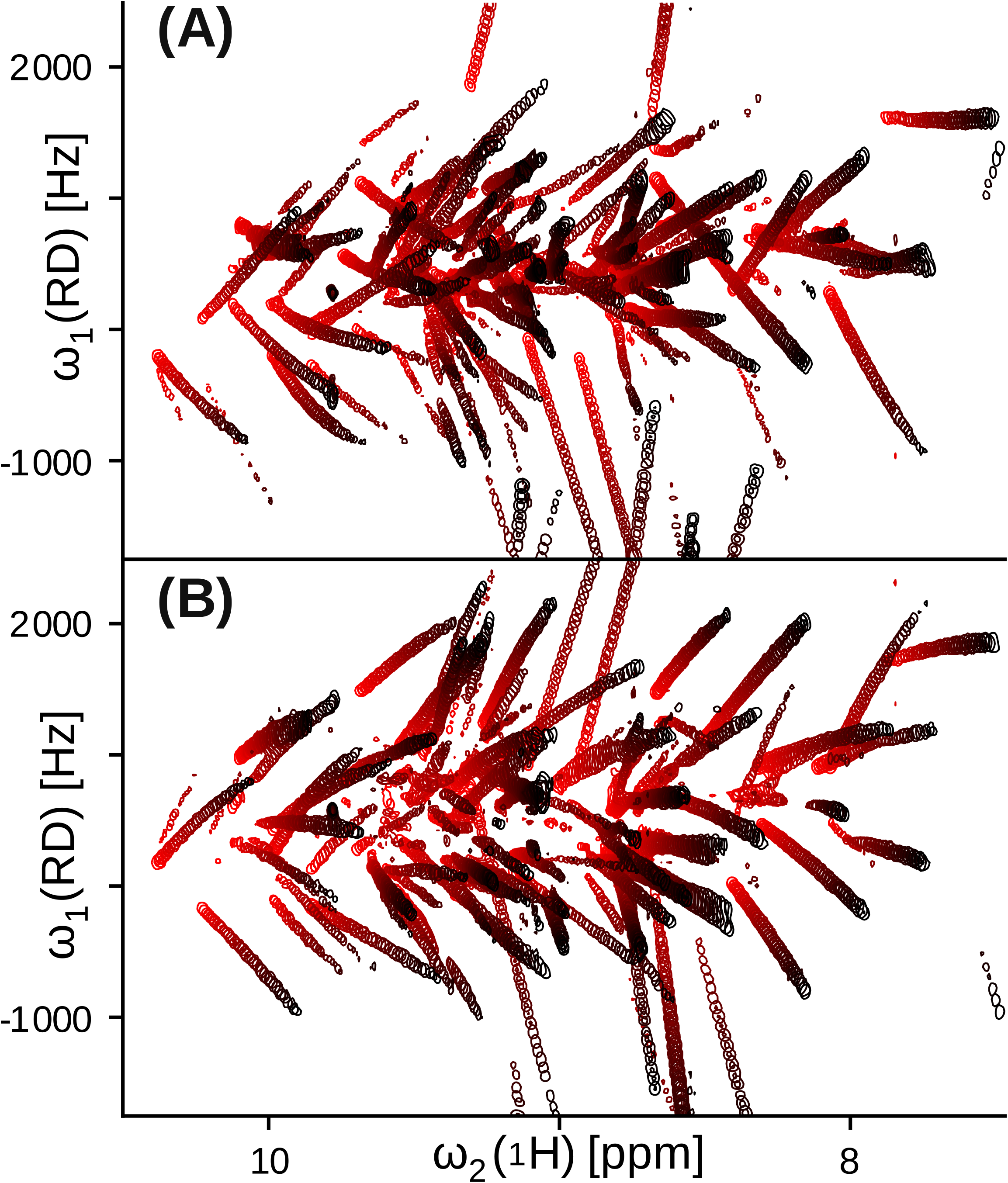
Variable-temperature RD-HSQC spectra of human ubiquitin (279– 318 K) with linearly modulated spectral widths. Forty contour plots are overlaid for each subspectrum panel, with temperature encoded by a continuous color gradient from blue (279 K, frame 1) to red (318 K, frame 40). **(a)** S□ and **(b)** S□ spectral plane overlays, each containing one cross-peak per residue. The S□ and S□ components are complementary spin-state-selective channels produced by the echo-antiecho editing of the RD-HSQC pulse sequence: at each temperature step, the S□/S□ peak pair for residue *k* is split symmetrically about the position of the conventional ^1^□N–^1^H HSQC cross-peak, with splitting magnitude proportional to the residue’s ^13^Cα rotating-frame offset according to |Δ□□□_RD(n)| = 2β(n)|Δ*v*□□□_C(n)| (Eq. 26), where β(n) = SWH_N(n)/SWH_C(n) is the frame-dependent mixing coefficient. Paired S□/S□ trajectories exhibit mirror-image motion in the mixed ^1^□N (RD) dimension, and their sum and difference recover the original per-residue ^1^□N and ^13^Cα chemical shift trajectories via the closed-form inversion Eqs. 28–30, confirming that the RD encoding is lossless when β ≠ 0.

Streak length and orientation in each panel encode the magnitude and direction of each residue^'^s net drift across the 40 K ramp. Compact, nearly circular spots indicate residues whose amide and ^13^Cα chemical shifts are weakly temperature-sensitive, consistent with deeply buried, strongly hydrogen-bonded backbone positions. Elongated streaks indicate residues whose amide shifts are strongly modulated by temperature, consistent with solvent-exposed or weakly hydrogen-bonded positions where backbone dynamics are sensitive to thermal perturbation. Peak motion with temperature originates primarily from changes in hydrogen bonding geometry at each backbone amide site, which modulates the ^1^H and ^1^□N chemical shifts at residue-specific rates governed by the drift velocity matrix **V** (Eq. 4 of the master equation).

The observed net drift direction incorporates a superposition of two contributions. The intrinsic contribution reflects genuine temperature-dependent changes in hydrogen bonding geometry, which typically drive backbone amide ^1^H and ^1^□N shifts upfield as temperature increases. Superimposed on this is a deuterium lock artifact: as the HDO resonance shifts upfield with increasing temperature, the spectrometer^'^s field-frequency lock compensates by adjusting B_0_ upward to return the deuterium lock signal to its reference position, inadvertently shifting all observed resonances downfield. The apparent peak velocities visible in the overlaid trajectories therefore reflect a superposition of intrinsic upfield drift from hydrogen bond weakening and a lock-induced downfield displacement common to all residues — a systematic effect that must be corrected before per-residue temperature coefficients can be interpreted in terms of hydrogen bond geometry. This controlled trajectory design, achieved through spectral-width modulation, transforms the variable-temperature series from a collection of discrete, independently analyzed observations into a continuous, physically trackable motion field amenable to video-processing-based analysis.

### Spectral-width modulation as a design variable

Given the master equation, the acquisition schedule—the sequence of SWH_C_(n) and ^13^C_α_ carrier offset 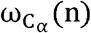 values across temperature steps—becomes a design variable. Modulating SWH_C_(n) varies the mixing coefficient β(n) = SWH_N_(n) /SWH_C_(n) and thereby the ^13^C_α_-driven S^+^/S^−^ splitting at each frame. We parameterise SWH_C_ (n) as a low-order polynomial in frame index:

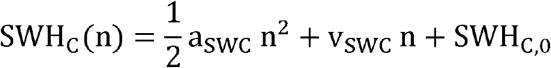

subject to a minimum guard SWH_C_(n) ≥ SWH_C,min_ to avoid aliasing of ^13^C_α_ resonances. Because β(n) = SWH_N_/SWHS_C_(n) and SWHS_N_ is held fixed, Eq. (8) uniquely determines the β (n) trajectory. An equivalent formulation holds if SWH_N_ is modulated instead; both descriptions are physically interchangeable. Only modulation of β(n), and thereby the relative displacement of S^+^ and S^−^ peaks, genuinely alters pairwise separability in Hz-space; carrier drift, being rank-1, cannot change pairwise separability between residues. Varying β converts the static ^13^C-driven splitting into a time-varying differential displacement that grows or contracts frame by frame, with the largest effect on residues with the largest 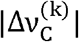.

### Cross-peak model-independent video super-resolution

Each temperature step shifts peaks by a small amount in frequency space. Across the series, the same peak is observed at many slightly different positions, providing more localization information than any single frame. We adapt the BasicVSR architecture (12, 13) to propagate spectral features bidirectionally across temperature steps—from lower temperatures forward and from higher temperatures backward—so that each frame is informed by resolved observations from both directions. Alignment is guided by the cross-peak intensity distribution, which plays the role of the optical flow field in natural video. The analogy holds because peak positions vary slowly and approximately linearly with temperature, producing a smooth, well-conditioned flow field amenable to standard optical-flow estimation. The approach makes no assumptions about drift linearity, residue-specific rates, or trajectory continuity, and is therefore robust to pathological cases such as conformational exchange broadening or cis/trans proline isomerization where a physical model would produce discontinuous or multistate trajectories.

BasicVSR++ (14) offers enhanced propagation and was therefore used. Figure 2 illustrates the improvement: reconstructed contours sharpen peak profiles relative to raw contours, with no systematic displacement of centroids. The processing pipeline applied to produce the blue (reconstructed) spectra operates in two sequential stages. In the first stage, the raw RD-HSQC frames are loaded into a spatio-temporal contrast-based spectral reconstruction (STSR) module. This step does not increase the spatial (spectral) resolution beyond the digital resolution ceiling set by the acquisition spectral width and number of time-domain points; rather, it stabilizes intensity across the temperature sequence by applying multiplicative 95th-percentile scaling with additive mean alignment to remove frame-to-frame gain variation, suppresses noise through temporal averaging of correlated spectral features, and eliminates baseline fluctuations, ensuring that each frame entering the super-resolution stage has a consistent, well-conditioned intensity profile. In the second stage, the normalised frame sequence is processed by a BasicVSR video super-resolution model trained on the Vimeo-90K dataset — a corpus of high-resolution natural video with smooth inter-frame motion. BasicVSR propagates spectral features bidirectionally across temperature steps (from lower temperatures forward and from higher temperatures backward through the series), aligning and fusing information from resolved frames to sharpen per-frame peak profiles at frames where peaks are close or overlapping. The output has four times the spatial resolution of the input frame sequence.

**Figure 2.**
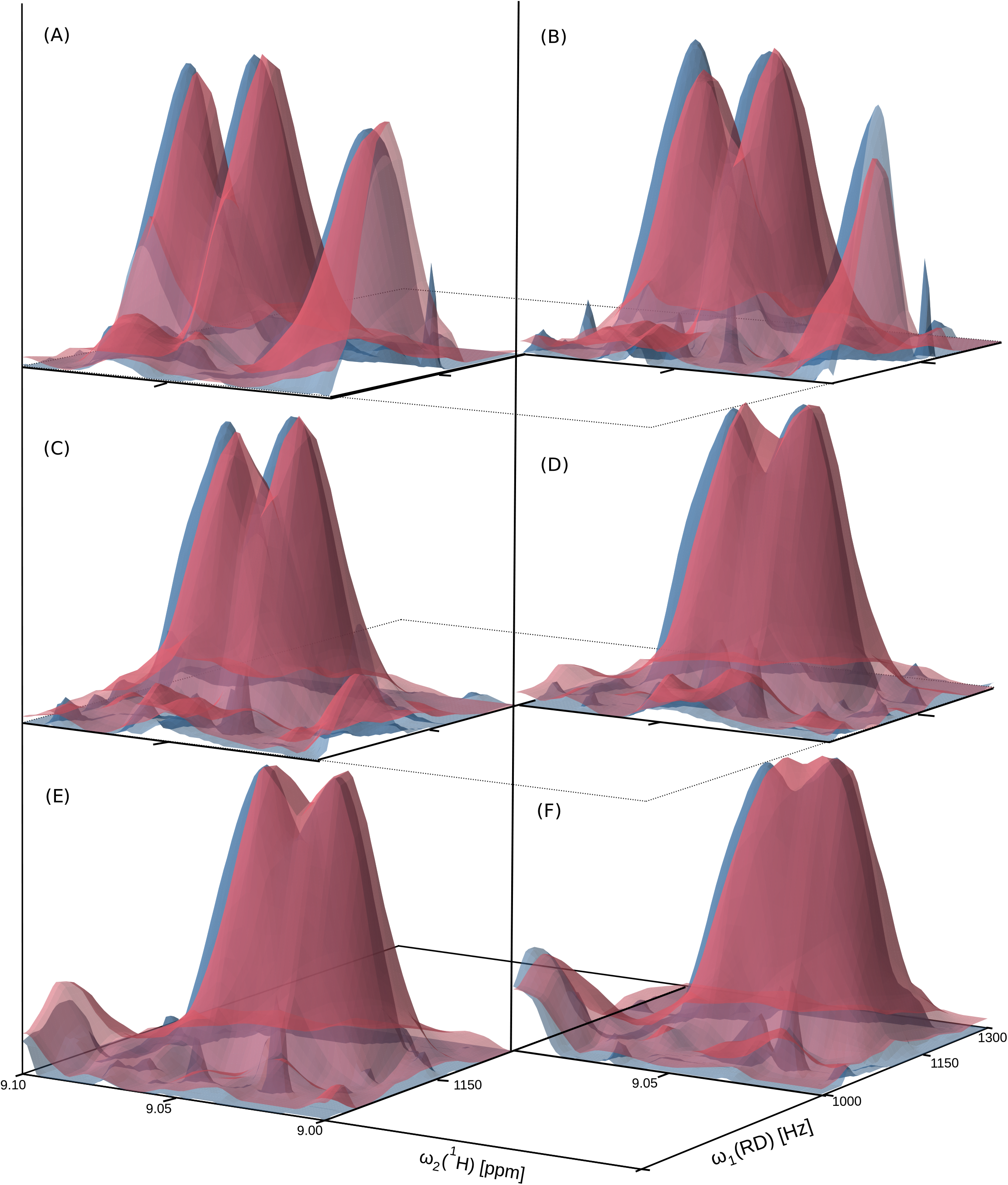
Peak model-independent video super-resolution reconstruction of RD-HSQC spectra of human ubiquitin. Panels **(a)–(f)** each show a cropped region of interest from the 2D RD-HSQC spectral window as a two-spectrum overlay: the raw spectrum (red contours), as processed directly in the Bruker TopSpin v4.05 following standard phase correction and baseline correction, is overlaid with the reconstructed spectrum (blue contours) produced by the full spatio-temporal processing pipeline. The six panels sample a descending temperature sequence from **(a)** 300 K to **(f)** 294 K in 1 K steps, so that the progression from panel **(a)** to **(f)** corresponds to cooling and the associated upfield drift of backbone amide resonances brings a pair of initially resolved cross-peaks progressively closer together and into partial overlap.

The region of interest selected for display contains two cross-peaks whose trajectories converge with decreasing temperature (Fig. 2). From panel (a) to (f), the groove — the intensity minimum between the two peaks in both the direct (^1^H) and indirect (RD ^1^□N) dimensions — progressively narrows as the peaks approach one another. In the raw spectra (red), by panel (f) the two peaks have merged into a single, unresolved contour, consistent with their separation falling below the digital resolution set by the acquisition parameters. In the reconstructed spectra (blue), the two peaks remain individually distinguishable as separate contours in panel (f), with a preserved intensity minimum between them, demonstrating that the temporal information pooled across the temperature series by the BasicVSR propagation step recovers per-frame resolution that is lost in single-frame analysis of the raw data. This resolution recovery carries no peak displacement: the centroids of well-resolved blue peaks are coincident with their red counterparts in the panels where both are resolved, confirming that the reconstruction introduces no systematic bias into downstream peak position estimates. The pipeline thus constitutes an effective preprocessing method for cross-peak tracking across variable-temperature RD-HSQC series, improving the fidelity of trajectory recovery in regions of the spectral window where resonances transiently overlap.

The cross-peak model-independent and model-based strategies are complementary in depth. A peak consistently occluded throughout the series cannot be recovered by model-independent methods alone; trajectory construction is indispensable in that case. Conversely, agnostic smoothing provides a quality floor that benefits all downstream stages regardless of acquisition quality. xxx

### Peak Tracking and Frequency Recovery

Per-frame peak detections—obtained by watershed segmentation (15) of the 2D intensity surface—are linked into continuous trajectories by a constant-velocity Kalman filter (16) with state vector (x,y,v_x_,v_y_), where x and y denote the ^1^H and apparent ^15^N frequencies respectively. The constant-acceleration prior is appropriate because backbone amide group spins shift temperature coefficients are approximately constant over moderate temperature ranges (4), producing trajectories that are nearly linear or slightly curved in frame index. A higher-order dynamical model would generate systematic prediction errors in the initial frames before the velocity state is well estimated, causing data-association failures precisely where they are most costly. The filter predicts each peak’s next position and updates upon detection; bidirectional initialization reduces track fragmentation at series endpoints.

The process noise covariance **Q** and measurement noise covariance **R** are tuned from the empirical distribution of per-step peak displacements in a held-out calibration subset of the ubiquitin dataset. For each accepted detection, the innovation (residual between predicted and measured position) is retained as a diagnostic: large, persistent innovations flag residues with non-linear temperature dependence, which are candidates for higher-order modelling or flagging as exchange-broadened.

#### Graph-based S^+^/S^−^ pairing and frequency recovery

Trajectory fragments from the S and S^−^ channels are first consolidated using a union-find data structure (17, 18) to group frame-level detections that belong to the same continuous trajectory. Pairing across channels exploits the constraint from Eq. (1): candidate S^+^ /S^−^ pairs are scored by the smoothness of their sum trajectory η^(k),+^(n) +η^(k),−^(n), which should follow a smooth, near-linear temperature dependence consistent with a pure ^15^N chemical shift trajectory. Pairings satisfying this smoothness criterion above a confidence threshold are retained as high-confidence assignments and used to bootstrap recovery of three-nucleus chemical shift trajectories via Eqs. (2) and (3). The recovery is lossless when β ≠0

In the reported series (Fig. 1) the 13C spectral width SWH_C_(n) and ^15^N spectral width SWH_N_(n) were modulated as functions of the frame index n across the 40-step temperature ramp (279–318 K, 1 K increments). Modulation was subject to the reciprocal-sum constraint 1/ SWH_N_(n) + 1/ SWH_C_(n) = C_const_(Eq. 8). This constraint maintained consistent digital resolution while enabling monotonic variation of the mixing coefficient β(n) throughout the series. All expected trajectories were successfully recovered from the dataset. Figure 6 reports reconstructed cross-peak trajectories encoding the detailed temperature dependencies of all backbone resonances in the test protein.

### Construction of optimal VT RD-HSQC Peak Trajectories

To study optimal construction of peak trajectories in RD-HSQC experiments, we simulated the evolution of cross-peak positions in temperature-resolved ^1^H–^15^N HSQC spectra of proteins with variable controls. Simulations were performed for temperature values ranging from 279 K to 315 K with a step of 1 K. At each temperature point the spectral coordinates of all peaks were calculated using backbone chemical shifts obtained from BMRB data. In these simulations each residue generates a pair of peaks whose coordinates evolve as temperature and controls change. Figure 5 shows the trajectories of these peaks through a sequence of temperature reprsented as moving points in a two–dimensional spectral plane expressed in frequency units (Hz). The detailed description of the simulations including temperature effect and trajectory controls is described in the Supplementary Information (section 1).

We formalize trajectory quality as a scalar scoring function over the space of acquisition schedules. Three criteria contribute: (i) repulsion—mean nearest-neighbour separation across the series, computed in the 2D observed spectral plane(η^±^, Δ*v*_H_); (ii) step displacement—frame-to-frame displacement variance, penalising schedules that produce large sudden jumps that would break tracker continuity; and (iii) boundary compliance—fraction of trajectory points within the spectral window, penalising configurations where peaks alias outside the acquired region and (iv) trajectories collision resolution. These criteria are combined into a single differentiable objective evaluated on simulated trajectories generated from Eq. (6) with chemical shifts statistics drawn from protein sequence databases (4).

This formalization permits principled optimization of the trajectories controls and, critically, enables agentic control of the design loop: the scoring function defines what constitutes a good acquisition schedule, and an autonomous agent traverses the low-dimensional parameter space (a_SWC_, v_SWC_, SWH_C,0_, and 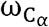) to maximise it. Example optimised trajectories are shown in Figure 5, with penalty components evaluated throughout the temperature sweep.

### Novel RDL-TROSY experiment

We developed a reduced-dimensionality longitudinal relaxation optimized experiment, denoted RDL-TROSY (Fig. 3), to maximize spectral resolution and controllability of cross-peak trajectories in temperature-resolved 2D [^1^H,^1^ □N] correlation spectra. In this sequence, excitation is restricted to amide protons or water and aliphatic protons using band-selective ^1^H pulses, enhancing longitudinal recovery of ^1^H^N^ polarization. The key design feature is that the entire period during which ^1^□N magnetization resides in the transverse plane is utilized for chemical shift encoding, thereby achieving maximal effective evolution time and, consequently, the highest attainable resolution in the indirectly detected dimension. Reduced-dimensionality encoding is implemented through controlled transfer between ^1^□N and ^13^Cα spins, producing the characteristic S(+) and S(–) components. Importantly, the sequence incorporates weak selective RF irradiation on the ^13^C channel during the ^1^□N→^13^C transfer period, or simultaneously on both ^13^C and ^1^□N channels, enabling targeted attenuation of specific cross-peaks. This feature allows dynamic suppression of selected resonances at defined time points, providing an experimental handle to resolve trajectory collisions and cross-peak occlusions in a temperature series.

**Figure 3:**
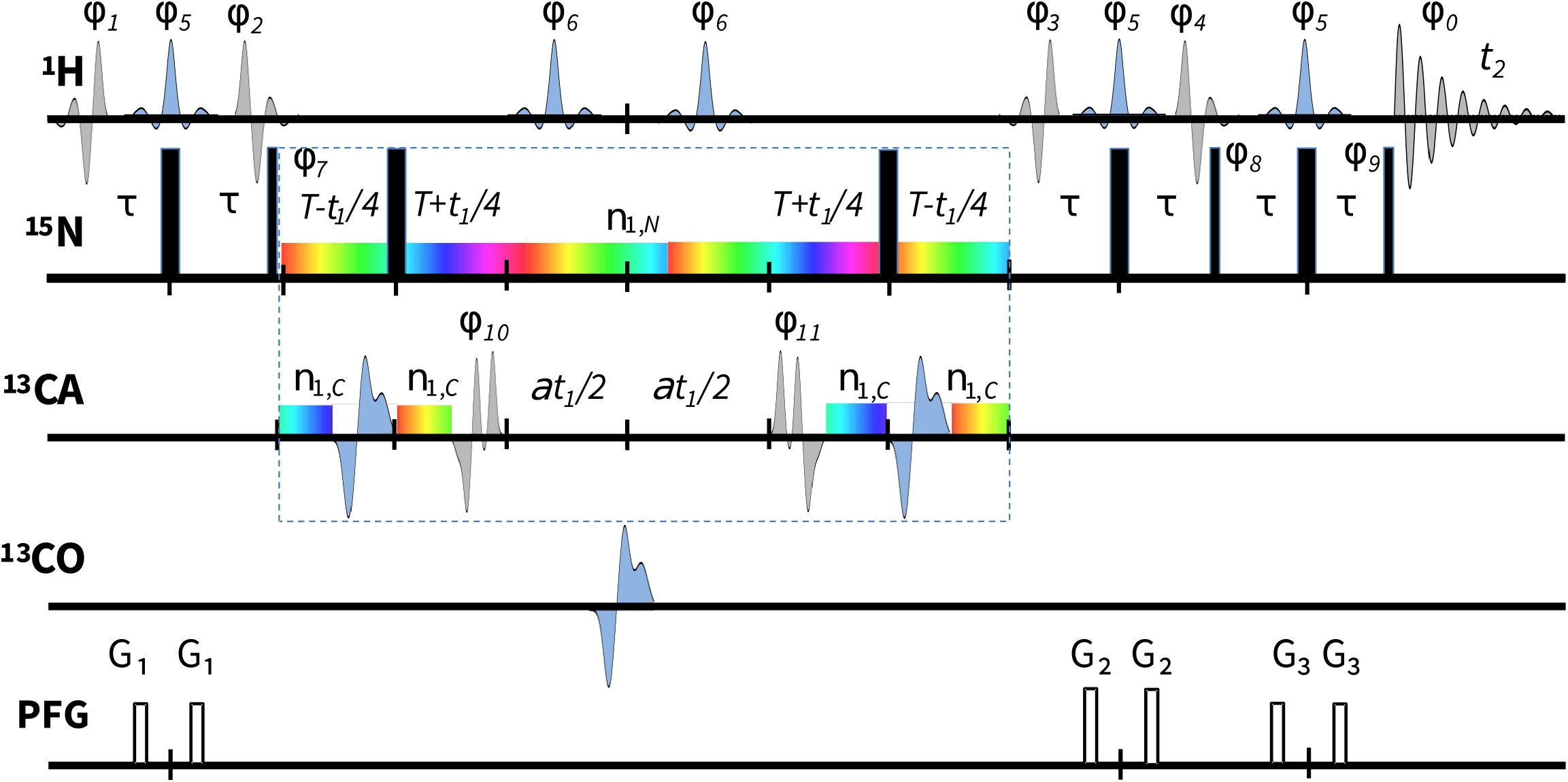
Experimental scheme for the Reduced-Dimensionality Longitudinal ^1^H relaxation optimized 2D [^1^H,^15^N]-TROSY (RDL-TROSY). The radio frequency pulses on ^1^H, ^15^N, ^13^C_α_, ^13^CO are applied at 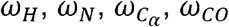 of 4.7, 118, 55, 174 ppm, respectively. Narrow and wide black bars indicate nonselective 90° and 180° pulses, respectively. The ^1^H shaped pulses are: *ϕ*_1-4_, ^1^H_*N*_ 5.5–10.0 ppm band-selective 1.5 ms excitation E-Burp2 pulses with the phases {*x*}, {y}, {y,y,y,y, −y, −y, −y, −y} and {*x*}, respectively. The shapes of *ϕ*_2,4_ are time reversed of *ϕ*_1,3_. The *ϕ*_5_ and *ϕ*_6_. are ^1^H_*N*_ 5.5–10.0 ppm and 5.5–1.0 ppm band-selective 1.8 ms refocusing Re-Burp pulses with the phase {*x*}. The ^13^C shaped pulses are 1.3 ms Gaussian cascade Q3 (dark shapes) and Q5 (light shapes) with the phases *ϕ*_10_ = {−*x*(8), *x*(8)}, *ϕ*_11 =_ {−*x*(8), *x*(8)} and {−*y*(8), *x*(8)} for the reduced-dimensionality in-phase and antiphase spectra,*S*_*in*_ and *S*_*anti*_, respectively. The spectra *S*(+) and *S*(−), are reconstructed by adding and substracting *S*_*in*_ and *S*_*anti*_, respectively. The phases *ϕ*_0,7−9_ are {*x*,−*x*,−*y,y,x*,−*x,y*,−*y*}, {*y*,−*y*,−*x,x*}, {*y*}, {*x*(4), −*x*(4)}, respectively. The echo-anti-echo transverse relaxation optimised spectroscopy (TROSY) phase discrimination pattern is applied to *ϕ*5 and *ϕ*_0_ for each *S*_*in*_ and *S*_*anti*_. The delays are *t* = 2.7 ms and *T* = 30 ms. The pulsed field gradients (PFGs) are G1: 80 G/cm; G2: 95 G/cm; G3: 70 G/cm. The colored shapes represent narrow band-selective phase-modulated radio-frequency irradiation with the radio-frequency (RF) field strength of *v*_1,*N*_ and *v*_1,*C*,_ with the corresponding offsets defined in the text. To generate cross-peak trajectories, the 2D RDL *S*(+), *S*(−)and, planes are measured with the temperature range (279–315 K) and variable settings of the ^13^C carrier frequency 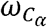, and the corresponding spectral width *SWH*_*N*_ and *SWH*_*C*_. The dashed box in the middle indicates the part of the pulse sequence used in the numeric simulations of the propagation of the density operator of two *J* - coupled spins ^13^C and ^15^N under the effects of the static Hamiltonian and the phase modulated weak RF irradiation

In contrast to conventional RD-HSQC implementations derived from folding the ^13^Cα dimension of a 3D HNCA experiment into the ^1^□N dimension, the RDL-TROSY sequence is specifically optimized for longitudinal relaxation and maximal encoding efficiency. Standard RD-HSQC inherits limitations from the parent 3D experiment, including fragmented evolution periods and reduced effective evolution time for ^1^□N, which constrain achievable resolution and lead to less controllable trajectory geometry. Furthermore, conventional approaches lack mechanisms for selective, time-resolved attenuation of individual peaks, making them vulnerable to peak overlap during trajectory evolution. By combining full-length ^1^□N evolution, TROSY-based relaxation optimization, and programmable weak selective irradiation, RDL-TROSY enables both improved spectral resolution and active manipulation of cross-peak trajectories. This enhanced control is essential for avoiding trajectory collisions and for enabling downstream computational strategies, such as reinforcement learning–guided optimization of acquisition parameters.

### Selective control of the trajectories at the collision point

The ability of the RDL-TROSY experiment to selectively attenuate individual cross-peaks in the S^+^ and S^−^ spectra is demonstrated in Figure 4. Panel (A) shows an overlay of two S^+^ spectra acquired with near zero (red) and maximal (black) weak RF irradiation applied on the ^13^C channel at a carrier position centered at 60 ppm, corresponding to the Cα chemical shift of Ile31. Under these conditions, a single targeted cross-peak is efficiently and selectively suppressed, while the remainder of the spectrum remains largely unaffected. This highlights the high residue specificity of the weak RF irradiation scheme, which exploits the narrow-band excitation profile to address individual resonances without perturbing neighboring peaks. The absence of noticeable distortions in surrounding signals confirms that the applied RF field of 28 Hz is sufficiently weak to avoid global perturbation while still achieving complete attenuation of the selected resonance.

**Figure 4:**
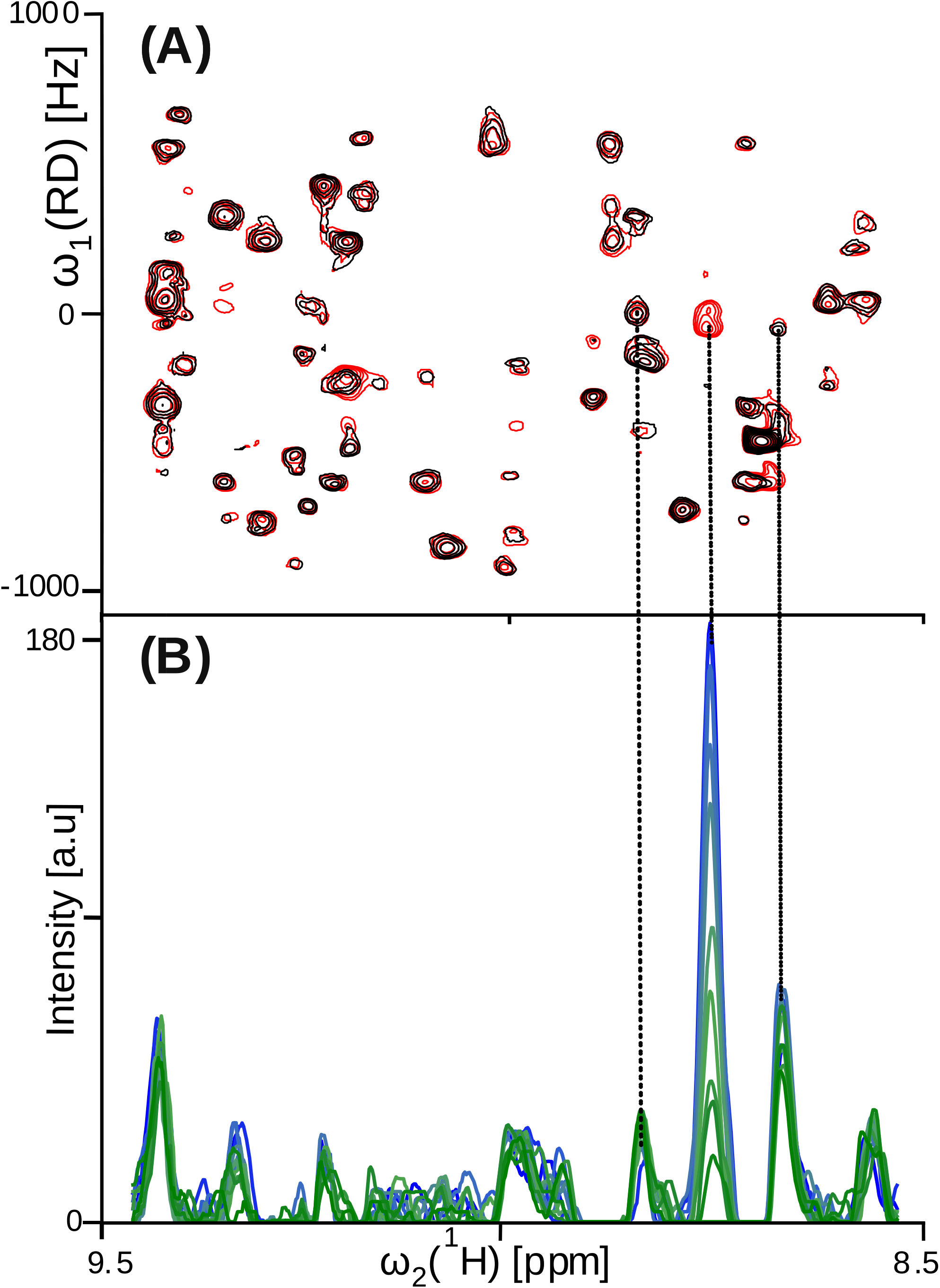
Selective and tunable cross-peak attenuation in RD-HSQC. **(a)** Overlay of two S spectra acquired with maximum (black) and minimum (red) RF irradiation at a targeted ^13^C offset. Only one peak (arrowed) is completely suppressed at using weak ^13^C radio-frequency irradiation *v*_1,*C*_ = 28 Hz centered at 60 ppm (CA chemical shift of Ile31), demonstrating residue-specific selectivity. **(b)** 1D slices through the suppressed peak (center) and two reference peaks (flanking) across the power series *v*_1,*C*_ = {0,□3.5,□5,□7,□11,□16,□22,□28,□40,□56,□80,□113,□143} Hz (colored from blue to yellow in the corresponding order), showing progressive attenuation of the target peak intensity as power increases.

**Figure 5:**
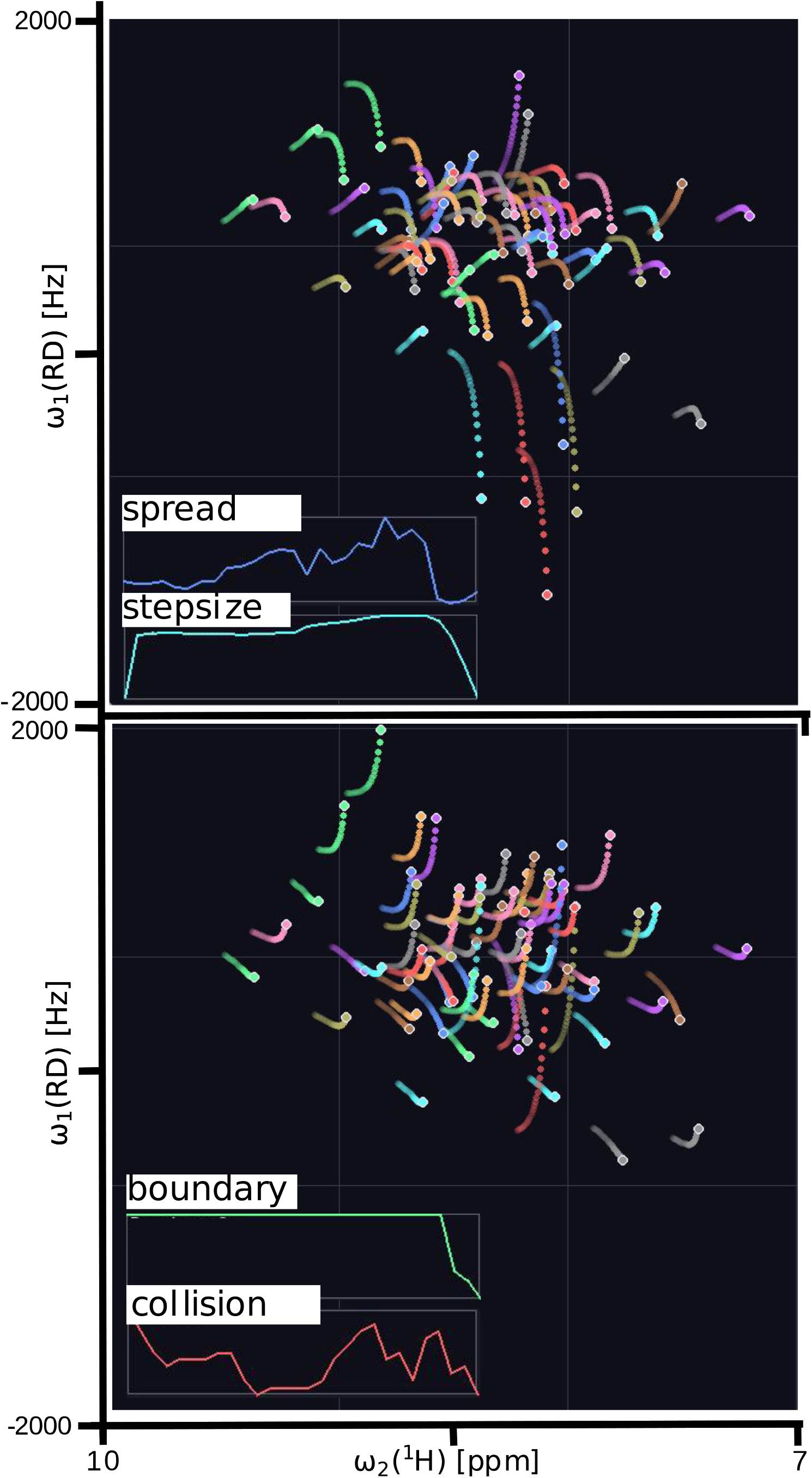
RD-HSQC peak trajectories generated using near-optimal control parameters. Each colored curve represents the temperature-dependent trajectory of a backbone amide resonance of ubiquitin (76 residues) across the simulated temperature range (279–315 K). The RD encoding produces two spectral planes (top and bottom panels), corresponding to the *S*(+) and *S*(−) components arising from folding of the ^13^C_α_ chemical shift into the indirect dimension. The trajectories are actively reshaped through global experimental controls acting on the ^13^C carrier frequency (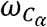) and the ^13^C spectral width (*SWH*_*C*_), whose rates and accelerations are dynamically updated with temperature while maintaining the reciprocal-sum constraint 1 / *SWH*_*N*_(n) + 1 / *SWH*_*C*_(n) + *C*_*const*_ (Eq. 8). The control parameters used in this example were 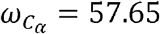 ppm with rate 0.39 ppm/K and acceleration 0.0187 ppm/K^2^, and *SWH*_*C =*_ 2617.8 Hz with rate 196.8 Hz/K and acceleration 9.86 Hz/K^2^. Insets show the evolution of representative penalty components used to evaluate trajectory optimality during the temperature sweep, including spread, step-size, boundary, and collision terms. These penalty functions quantify peak separation, smoothness of motion, distance from spectral boundaries, and avoidance of peak overlaps, respectively, and together define the objective function used for trajectory optimization.

**Figure 6:**
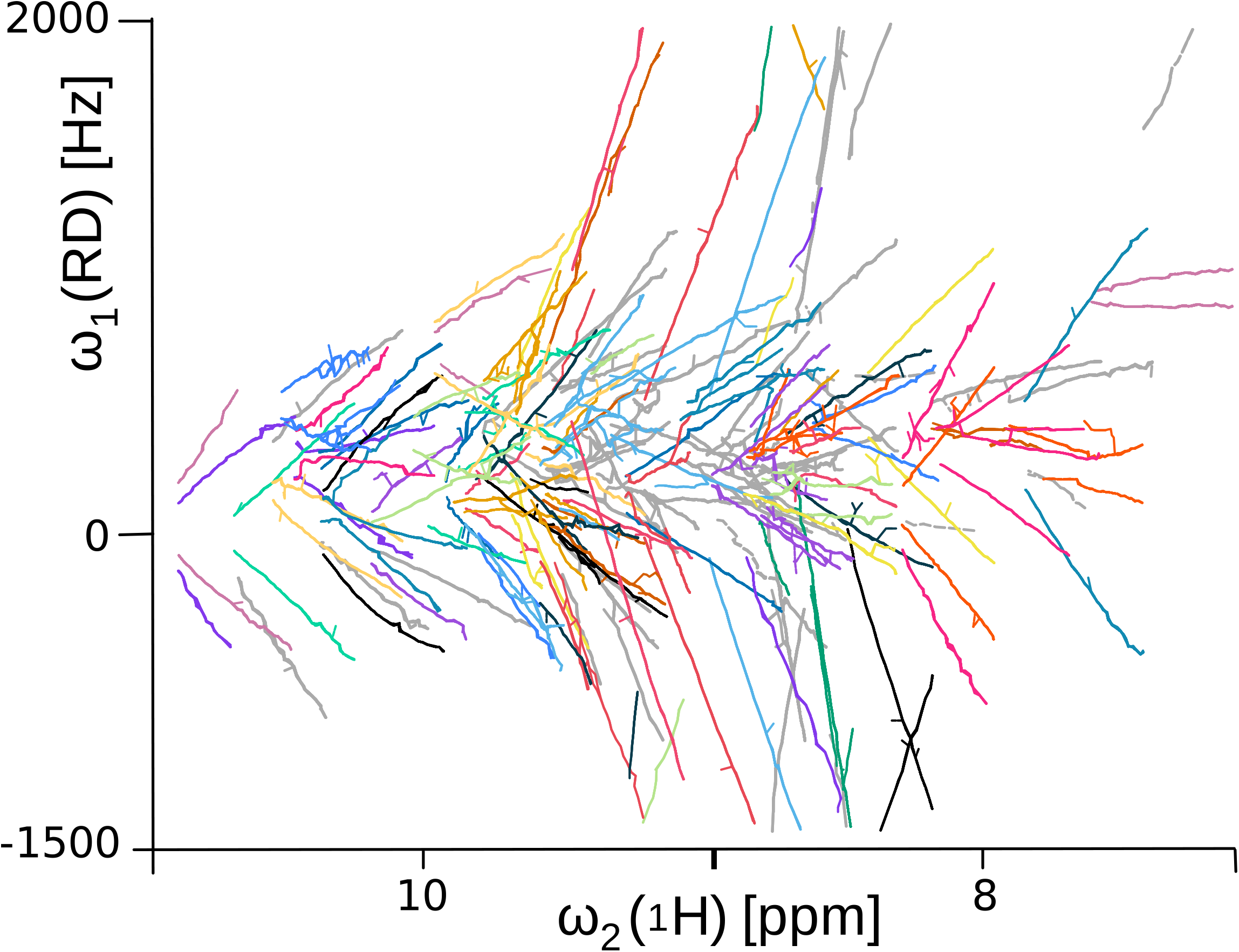
Automated S+/S− trajectory pairing via iterative RMSD bootstrapping. Each colored trajectory represents a matched S+/S− trajectory pair, with paired trajectory sharing the same color. Grey trajectories in the background indicate unmatched candidate islands. Pairing was performed by iterative Hungarian assignment with a progressively relaxed RMSD threshold (starting from 0, incremented by 0.001 per round over 20,000 iterations). The RMSD serves as the sole matching criterion, exploiting the physical constraint that the ^1^H chemical shift drift (Δ*v*_H) is identical for the S+ and S− peaks of the same residue, producing coincident horizontal trajectories across the temperature series.

Figure 4B further quantifies this effect by showing one-dimensional slices through the suppressed peak and two neighboring reference peaks across a series of increasing RF irradiation strengths (0–143 Hz). The intensity of the targeted peak decreases smoothly and monotonically with increasing RF power, ultimately reaching near-complete suppression, whereas the flanking peaks remain largely unchanged across the entire power range. This continuous and controllable attenuation demonstrates that weak RF irradiation can be used as a finely tunable parameter to modulate peak intensities in a predictable manner. The attenuation effect of the weak rf-irradiation was extensively numerically modeled (Supplementary material, section 4) confirming experimentally observed attenuation levels (Fig. 3S). Interestingly, our simulations show that to almost complete and selective peak intensity attenuation can be achieved with a combined weak ^15^N and ^13^C irradiation of only 5 to-10 Hz in the field strength (Fig. 4S) creating a very precise selection of the targeted trajectory.

Importantly, this introduces a new experimental control dimension for trajectory optimization: selective, time-resolved masking of specific resonances. In the context of AI-driven trajectory construction, such weak RF interventions can be formulated as discrete, low-cost actions that temporarily suppress colliding peaks, enabling the agent to resolve trajectory overlaps without altering the underlying spectral encoding. This capability provides a direct experimental counterpart to algorithmic “masking” operations and significantly enhances the feasibility of agentic control over peak trajectories in complex spectra.

## Discussion

In this work, we introduce a conceptual and experimental framework that redefines variable-temperature NMR as a spatiotemporal problem in which cross-peaks evolve along a controllable trajectory axis. Rather than treating temperature as a perturbation to be minimized or corrected, we exploit it as a structured, physically meaningful variable that encodes additional information into the spectral domain. In combination with reduced-dimensionality encoding, this transforms a series of 2D spectra into a trajectory-resolved dataset in which otherwise overlapping resonances can be separated through their differential motion. In contrast to conventional multidimensional NMR, where additional spectral resolution is obtained by extending Fourier dimensions at the cost of acquisition time, our approach redistributes dimensionality into a controllable external variable, enabling efficient extraction of multi-nuclear chemical shift information within a comparable experimental time.

This framework provides both an experimental and computational advantage. On the experimental side, modulation of the RD mixing parameter and spectral widths allows active steering of cross-peak trajectories, converting latent ^13^C chemical shift diversity into time-dependent separability. The RDL-TROSY experiment further maximizes the achievable resolution by fully utilizing the ^1^□N transverse evolution period and introduces a novel control dimension through weak selective RF irradiation. This selective attenuation acts as a precise, residue-specific intervention that resolves trajectory collisions without perturbing the global spectral encoding. On the computational side, the trajectory formulation enables direct application of computer vision methodologies, including video super-resolution and multi-object tracking, which leverage temporal redundancy to recover information lost in individual frames. Together, these elements establish a unified framework in which acquisition design and data analysis are intrinsically coupled.

A key implication of this work is that spectroscopic dimensionality need not be restricted to traditional frequency axes. External variables such as temperature, pH, ionic strength, or ligand concentration can be incorporated as controlled trajectory-generating dimensions, provided that their effects on chemical shifts are sufficiently systematic. This generalization suggests a broader paradigm of “trajectory-encoded spectroscopy,” in which the experiment is designed to produce maximally informative motion of resonances rather than static separation in high-dimensional frequency space. Within this paradigm, the information content arises not only from instantaneous peak positions but also from their evolution, effectively introducing derivative constraints that improve identifiability of overlapping signals.

The formulation of trajectory quality as a differentiable objective function further enables automated optimization of acquisition parameters. In this study, we demonstrate that global controls such as carrier frequency and spectral width can be tuned to maximize peak separability while maintaining smooth, trackable motion. Importantly, the addition of selective RF attenuation introduces a discrete control mechanism that can be used to resolve transient peak overlaps. This naturally lends itself to a reinforcement learning formulation, in which an agent iteratively adjusts acquisition parameters and selective interventions to maximize a global reward function. Unlike conventional automation, which executes predefined protocols, such an agentic framework directly influences the physical experiment in real time, enabling adaptive and protein-specific optimization of the acquisition process.

Several limitations should be noted. The approach relies on approximately linear temperature dependence of chemical shifts and requires that the protein remains structurally stable over the sampled temperature range. Strong conformational exchange, unfolding transitions, or non-linear drift may complicate trajectory modeling and tracking. In addition, the requirement for dense sampling across temperature introduces a trade-off between temporal resolution and total acquisition time, although this is partially mitigated by the use of efficient 2D experiments. Finally, while the current implementation focuses on backbone assignment, extension to side-chain correlations or distance restraints will require further development of both pulse sequences and trajectory models.

Overall, the present work establishes a bridge between NMR spectroscopy, trajectory-based signal encoding, and modern computer vision methodologies. By integrating experimental control, physical modeling, and data-driven analysis within a unified framework, it opens a pathway toward adaptive, agent-guided NMR experiments in which acquisition and interpretation are co-optimized. This perspective suggests that future developments in NMR may increasingly rely not only on improved hardware or pulse sequences, but also on intelligent control strategies that actively shape the information content of the experiment.

## Conclusions

We have described a framework that reconceptualizes the VT-NMR experiment as a video processing problem and demonstrated concrete consequences of that reconceptualization for cross-peak resolution and residue assignment. The temperature axis, conventionally treated as a complication, is recast as a pseudo-temporal axis along which inter-frame pooling resolves peaks irresolvable in any individual spectrum. The RD-HSQC is the natural vehicle: its explicit ^13^C_α_ encoding provides spectral, temporal, and topological discrimination simultaneously, and its exact mathematical form provides the basis for principled acquisition design.

The agnostic and model-based strategies are complementary: the former improves per-frame quality without physical assumptions; the latter converts latent ^13^C_α_ diversity into engineered inter-trajectory separation through spectral-width modulation. A formal scoring function, a selective intensity-control pulse sequence element, and the agentic design loop together define a path toward fully automated acquisition optimisation. The framework is protein-independent; its inputs are backbone chemical shift databases and the target primary sequence, and its output is an optimised acquisition schedule calibrated to the specific assignment problem at hand.

## Materials and Methods

Human ubiquitin ^15^N/^13^C uniformly labeled was purchased from Sigma-Aldrich (MilliporeSigma). NMR spectra were measured using Bruker Avance III NMR spectrometer operating at 700 MHz magnetic field strength. Specgra were referenced relative to the internal DSSThe RD-HSQC was acquired quadrature selective ^13^C_α_ editing, yielding separate S^+^ and S^−^ free induction decays (FIDs) by acquiring the in-phase (*S*_in_) and antiphase (*S*_anti_) spectra in an interleaved fashion (10, 11). The temperature ramp spanned 279–318 K in 1 K increments (40 steps); one complete RD-HSQC was acquired per step following 1 min thermal equilibration. Temperature calibration was performed using the standard methanol sample (2).

Raw Bruker TopSpin free induction decays (FIDs) were apodised (cosine-bell in t_1_, exponential in t_2_), zero-filled to twice the acquisition size in both dimensions, Fourier transformed, and phased. S^+^ and S^−^ subspectra were computed by addition and subtraction of the *S*_in_ and *S*_anti_ spectra. Both channels were assembled into video sequences at 4 fps with 1:1 array-to-pixel grayscale encoding and normalised by multiplicative 95th-percentile scaling with additive mean alignment. Inter-frame consistency was assessed by the structural similarity index measure (SSIM) (20). Agnostic super-resolution, per-frame peak detection, Kalman tracking, graph-based classification, and frequency-coordinate recovery were implemented in Python 3.10 with graphics processing unit (GPU) acceleration (CUDA 11.8).

## Acknowledgements

We thank Prof Vladislav Orekhov for fruitful early discussions and his advice for the project.

