## Supplementary Information for "NMR as a Video-(game): Constructing Super-Resolution Cross-peak Trajectories in Protein Spectroscopy"

### 1.Simulation of RD-HSQC Peak Trajectories

To study optimal construction of peak trajectories in reduced-dimensionality (RD) HSQC experiments, we simulated the evolution of cross-peak positions in temperature-resolved $^{1}$H–$^{15}$N HSQC spectra of proteins. Each residue contributes one backbone amide peak in a conventional HSQC experiment. In the RD-HSQC implementation, the peak is split into two components due to incorporation of the $^{13}$C$_{\alpha}$ chemical shift into the indirect dimension, producing two spectra denoted $S(+)$ and $S(-)$, where the $^{15}$N coordinate is shifted by $\pm\Delta/2$ depending on the encoded $^{13}$C$_{\alpha}$ chemical shift. Consequently, each residue generates a pair of peaks whose coordinates evolve as experimental parameters change. The trajectories of these peaks through a sequence of temperature points are treated as moving points in a two–dimensional spectral plane expressed in frequency units (Hz).

Simulations were performed for temperature values ranging from 279 K to 315 K with a step of 1 K. At each temperature point the spectral coordinates of all peaks were calculated using backbone chemical shifts obtained from BMRB data. The RD representation produces two independent spectral planes ($S(+)$ and $S(-)$) that are treated separately in the trajectory optimization.

#### *Temperature-induced variation of the chemical shifts*.

The parametric forms that suitable for backbone $^{1}$H$^{N}$, $^{15}$N, $^{13}$C'/$^{13}$Cα chemical shifts vs absolute temperature $T$(K) using a quadratic around a reference temperature $T_0$(e.g., 298.15 K):

$$\delta_X(T) = \delta_{X,0} + a_X\,(T - T_0) + b_X\,(T - T_0)^2$$

- $X \in \{^{1}\mathrm{H}^{N},\ ^{15}\mathrm{N},\ ^{13}\mathrm{C}'$ (or $^{13}\mathrm{C}\alpha)\}$
- $\delta_{X,0}$: shift at $T_0$
- $a_X$: linear temperature coefficient (ppm/K)
- $b_X$: curvature (ppm/K$^2$), usually small

For each individual simulation a random value of the typical parameter ranges (folded proteins, aqueous buffer) is selected:

**$^{1}$H$^{N}$**

- $a_H \approx -0.002$to $-0.015$ppm/K
  (most often $-0.004$to $-0.010$ppm/K; more negative = stronger H-bonding / solvent exposure changes)

- $b_H \approx -5 \times 10^{-5}$to $+5 \times 10^{-5}$ppm/K$^2$
  (often close to 0 over 5–45 °C)

**$^{15}$N**

- $a_N \approx -0.02$to $+0.02$ppm/K
  (often small magnitude; sign can vary by residue/secondary structure)
- $b_N \approx -5 \times 10^{-4}$to $+5 \times 10^{-4}$ppm/K$^2$

**$^{13}$C' (carbonyl)**

- $a_{C'} \approx +0.001$to $+0.01$ppm/K
  (often weakly positive)
- $b_{C'} \approx -1 \times 10^{-4}$to $+1 \times 10^{-4}$ppm/K$^2$

**$^{13}$Cα**

- $a_{C\alpha} \approx -0.002$to $+0.006$ppm/K
- $b_{C\alpha} \approx -1 \times 10^{-4}$to $+1 \times 10^{-4}$ppm/K$^2$

This quadratic is the best starting point for fitting and comparing residues.

#### 2.Control Parameters Governing Trajectory Geometry

The geometry of RD peak trajectories can be manipulated through acquisition parameters that determine the mapping between chemical shifts and the spectral frequency axes. Two global experimental controls were used to steer trajectories:

1. **$^{13}$C carrier frequency** $o_C$ (ppm). This parameter determines the reference frequency for $^{13}C_\alpha$ encoding and affects the relative displacement of peaks in the RD dimension.
2. **$^{13}$C spectral width** $SW_C$ (Hz). Changing the spectral width modifies the scaling of the RD encoding and therefore alters the direction and curvature of trajectories in the spectral plane.

Both parameters were modeled dynamically using second–order control equations analogous to a physical system with velocity and acceleration. For example, the evolution of the $^{13}$C carrier was described by

$$\begin{aligned} v_C(t+1) &= v_C(t) + a_C(t)\,\Delta T, \\ o_C(t+1) &= o_C(t) + v_C(t+1)\,\Delta T, \end{aligned}$$

where $v_C$ and $a_C$ denote the rate and acceleration of the carrier frequency with respect to temperature. A similar control law was used for the spectral width $SW_C$. These acceleration variables constitute the primary control actions used to shape the trajectories.

To maintain approximately constant spectral resolution across frames, the spectral widths in the $^{15}$N and $^{13}$C dimensions were constrained by

$$\frac{1}{SW_N} + \frac{1}{SW_C} = \text{constant}.$$

This relation ensures that the effective RD spectral resolution remains approximately invariant during trajectory optimization.

#### 3.Penalty Functions for Trajectory Optimality

Optimal trajectories are defined as those that remain trackable across temperature points, avoid peak overlap, and utilize the available spectral window efficiently. These objectives were implemented through a set of penalty functions that together form a global loss function.

##### 3.1.Repulsive Spread Loss

To discourage peak overlap, a repulsive energy term was introduced between peak pairs within each spectral plane. For peaks $i$ and $j$ separated by distance $d_{ij}$ (in Hz), the pairwise repulsive energy was defined as

$$E_{ij}^{\text{repulse}} = \frac{K_{\text{repulse}}}{(d_{ij} - d_{\text{clash}} + \epsilon)^2},$$

where $d_{\text{clash}}$ defines the minimum acceptable separation between peaks and $\epsilon$ prevents singularities. The interaction is evaluated only for peak pairs closer than a cutoff distance $d_{\text{cutoff}}$. The total spread loss is then

$$E_{\text{repulse}} = \frac{1}{M}\sum_{i<j} E_{ij}^{\text{repulse}},$$

where $M$ is the number of active peak pairs within the cutoff. The calculation is performed independently for the $S(+)$ and $S(-)$ spectra and the results are summed.

##### 3.2.Boundary Repulsion Loss

To prevent peaks from approaching the spectral boundaries, a similar repulsive energy was defined between each peak and the nearest spectral boundary:

$$E_i^{\text{bound}} = \frac{K_{\text{bound}}}{(d_i - d_{\text{bound}} + \epsilon)^2},$$

where $d_i$ is the distance to the nearest boundary and $d_{\mathrm{bound}}$ defines a safety margin. Only peaks within a cutoff distance $d_{\mathrm{bound\_cutoff}}$ from a boundary contribute to this term.

#### 3.3.Collision geometry and resolvability by video super-resolution

During temperature-dependent RD–HSQC experiments the cross-peaks follow continuous trajectories in the spectral plane. Occasionally two trajectories intersect, producing a temporary spectral overlap (collision). Whether such collisions can be resolved computationally depends on the local geometry of the intersecting trajectories and on the temporal information available from neighboring frames of the temperature series.

To analyze this effect, we treat each peak trajectory as a time-parametrized curve in the spectral plane and define the *collision angle* $\theta$ as the angle between the instantaneous tangent vectors of the two trajectories at the intersection point. The value of $\theta$ determines the degree to which the two peaks can be separated using video super-resolution (VSR) methods that exploit temporal information from neighboring frames.

When the collision angle is sufficiently large ($\theta > \theta_{\mathrm{thr}}$), the two trajectories approach the intersection point from clearly distinguishable directions. In this regime, frames preceding and following the collision contain spatially separated peaks whose trajectories can be extrapolated through the overlap region. Consequently, VSR algorithms can reconstruct the two signals and fully resolve the peaks at the collision point (Fig. X, blue occlusion point). In contrast, when the trajectories intersect at a small angle ($\theta < \theta_{\mathrm{thr}}$), the peaks approach the collision along nearly parallel directions. In this case the temporal evolution of the peaks provides insufficient geometric information to separate the two signals, and VSR reconstruction may fail.

The threshold angle $\theta_{\mathrm{thr}}$ depends on the effective point-spread function (PSF) of the NMR cross-peaks and on the number of available frames in the time series. In practice the PSF is dominated by the spectral linewidth (approximately the full width at half height), which defines the minimal spatial separation required for two peaks to be distinguishable. If the projected separation of the trajectories in adjacent frames remains larger than the PSF width, the two signals can be reconstructed reliably by VSR. Conversely, when the separation falls below this scale over several consecutive frames, the peaks become indistinguishable.

To incorporate this concept into the trajectory optimization framework, collisions are evaluated not only by the minimal distance between peaks but also by their collision angle. Intersections with $\theta > \theta_{\mathrm{thr}}$ are considered *resolvable collisions* and do not incur a penalty during trajectory optimization. Intersections with $\theta < \theta_{\mathrm{thr}}$ are classified as *sharp-angle collisions* and are penalized, because they cannot be reliably separated using VSR alone. In such cases an additional experimental intervention may be required, for example selective attenuation of one resonance using weak RF irradiation on the $^{13}$C or combined $^{13}$C/$^{15}$N channels.

In the reinforcement-learning framework used for trajectory optimization, this geometric criterion is incorporated into the loss function as an angle-dependent penalty. The agent therefore learns to construct trajectories that either avoid sharp-angle collisions or transform them into wide-angle intersections that remain resolvable by the VSR reconstruction procedure.

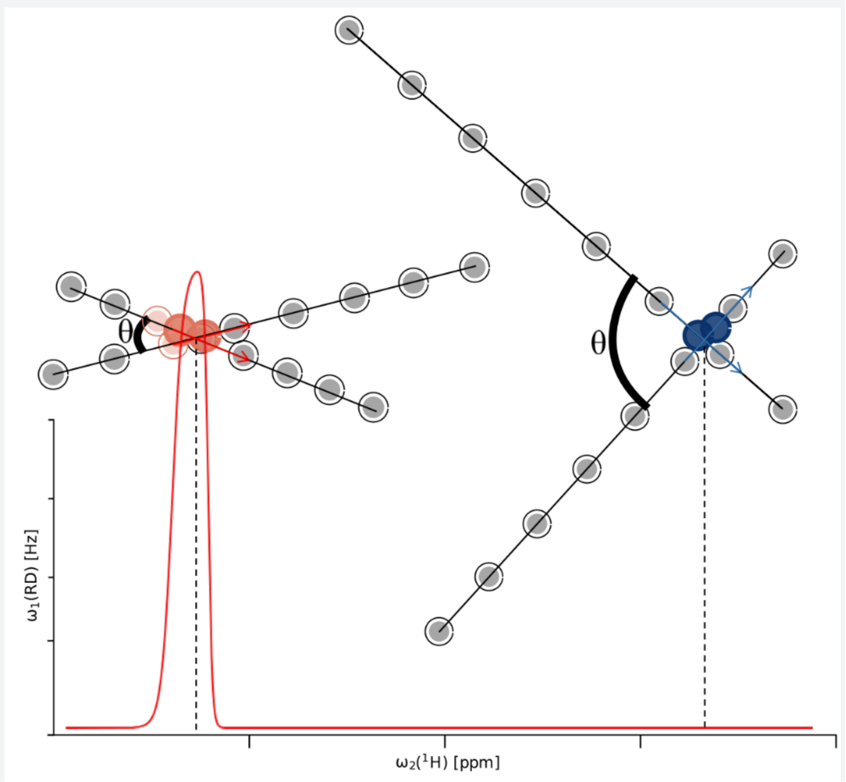


**Figure** 1S. Schematic illustration of two types of trajectory collisions in reduced-dimensionality spectra. When two peak trajectories intersect with a large collision angle Θ the temporal information contained in neighboring frames allows video super-resolution (VSR) reconstruction to resolve the peaks at the occlusion point. In contrast, collisions occurring at small angles (left, red) produce nearly parallel trajectories that remain indistinguishable within the point-spread function of the cross-peaks, primarily determined by their linewidth at half height. Such sharp-angle collisions are therefore penalized during optimal trajectory construction (red curve), whereas wide-angle intersections are not penalized. In cases where sharp collisions cannot be avoided, selective weak RF irradiation on $^{13}C$ or combined $^{15}N$ and $^{13}C$ channels can be applied to attenuate one of the peaks and restore resolvability.

##### 3.4. Step Size Regularization

To maintain smooth trajectories, a reward term favors moderate peak displacement between consecutive temperature frames. Motion that is too small leads to stagnation, while excessively large motion is penalized.

#### 4. Reinforcement Learning for Trajectory Construction

The trajectory optimization problem can be formulated as a reinforcement learning (RL) task. In this formulation the system state consists of the current spectral coordinates of all peaks, their recent trajectory history, and the current values of the experimental parameters. The agent actions correspond to discrete adjustments of the control accelerations $a_C$ and $a_{SW}$ that modify the carrier frequency and spectral width. The reward signal is derived from the negative of the loss function described above.

The RL agent interacts with the simulated environment over a sequence of temperature steps, learning a policy that maximizes cumulative reward while avoiding collisions and maintaining well-separated trajectories. Additional actions can optionally represent selective RF interventions that temporarily suppress specific peaks to resolve trajectory occlusions. Through repeated episodes and randomized chemical shift perturbations, the learned policy can generalize to proteins with different size and chemical shift distributions.

##### 4.1. Simulation of Reduced-Dimensionality 1D Spectra

Reduced-dimensionality 1D spectra were simulated using an in-house Python program implementing density-operator propagation for a coupled heteronuclear two-spin-1/2 system representing a $^{13}\mathrm{C}$–$^{15}\mathrm{N}$ spin pair. The spin dynamics were calculated in the rotating frame using explicit numerical propagation of the density operator

$$\rho(t+\Delta t) = e^{-iH\Delta t}\rho(t)e^{+iH\Delta t},$$

with a propagation step of $\Delta t = 1\ \mu s$.

The static Hamiltonian was defined as

$$H_0 = 2\pi(\Delta\omega_C C_z + \Delta\omega_N N_z + J_{CN} C_z N_z),$$

where the chemical shift offsets were set to $\Delta\omega_C = 100$ Hz and $\Delta\omega_N = 50$ Hz, and the scalar coupling constant was $J_{CN} = 15$ Hz.

Time-dependent radiofrequency irradiation was included through

$$H_{\mathrm{RF}}(t) = 2\pi[\omega_{1,C}(t)(\cos\phi_C(t)C_x + \sin\phi_C(t)C_y) + \omega_{1,N}(t)(\cos\phi_N(t)N_x + \sin\phi_N(t)N_y)].$$

To mimic irradiation with shifted excitation maxima, the RF phases were modulated linearly in time according to

$$\phi_C(t) = \phi_{C,0} + 2\pi\Delta\Omega_{\mathrm{RF},C}t, \qquad \phi_N(t) = \phi_{N,0} + 2\pi\Delta\Omega_{\mathrm{RF},N}t,$$

with phase-modulation offsets $\Delta\Omega_{\mathrm{RF},C} = 100\ \mathrm{Hz}$ and $\Delta\Omega_{\mathrm{RF},N} = 50\ \mathrm{Hz}$.

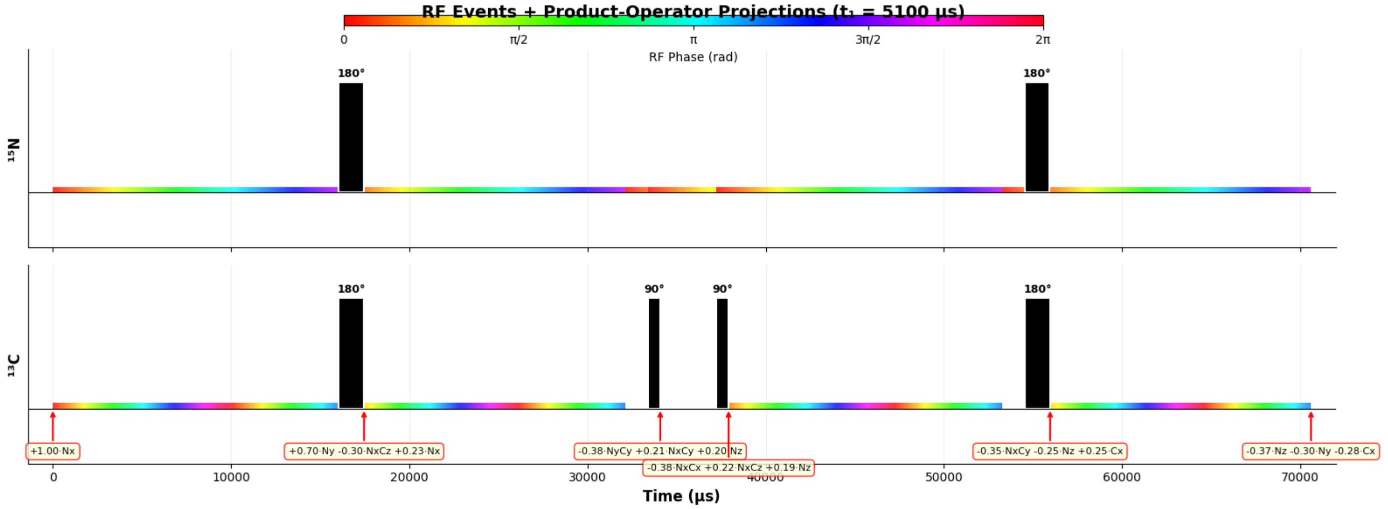


**Figure** 2S. The schematic diagram of the simulated part of the RDL-TROSY pulse sequence described in Figure 3 (main text) was used to simulate the propagation of the density operator of two J-coupled spins $^{13}$C and $^{15}$N under the effect of the static Hamiltonian and the phase-modulated weak RF irradiation. The colored shapes represent narrow band-selective phase-modulated radio-frequency irradiation with the rf-field strength of $\nu_{1,N}$ and $\nu_{1,C}$ with the corresponding offsets $\Delta\Omega_{\mathrm{RF},C}$ and $\Delta\Omega_{\mathrm{RF},N}$ in Hz. The pulse sequence is drawn to scale in the timeline corresponding to the $t_1$ evolution time of 5.1 ms. The density operator projections to the two-spin orthogonal basis at the time points specified by the red arrows are shown.

Simulations started from transverse $^{15}$N magnetization ($N_x$). Magnetization transfer between $^{15}$N and $^{13}$C spins was modeled using delays corresponding to the scalar-coupling transfer condition with $J_{CN} = 15$ Hz. During the transfer periods a weak continuous RF field was applied on $^{13}$C. The RF amplitude used during the evolution period was $\omega_{1,C} = 0$ Hz, while during transfer periods it was swept over

$$\omega_{1,C}^{(\mathrm{TE})} = \{0,\ 3.5,\ 5,\ 7,\ 11,\ 16,\ 22,\ 28,\ 40,\ 56,\ 80,\ 113,\ 143\}\ \mathrm{Hz}.$$

To generate reduced-dimensionality spectra, the phase of the first $^{13}$C 90° pulse was cycled according to

$$\phi_{90^\circ C}^{(1)} = \{0, \pi\}\ \mathrm{rad},$$

while the second $^{13}\mathrm{C}$ 90° pulse was applied with fixed phases

$$\phi^{(2)}_{90^\circ C} = \pi \ \mathrm{or} \frac{\pi}{2}$$

for $S_{in}$ or $S_{anti}$ spectra, respectively. The reduced-dimensionality evolution period was scaled relative to the indirect evolution time according to

$$t_{\mathrm{evol}} = \alpha \frac{t_1}{2},$$

with scaling coefficient $\alpha = 1.5$. The indirect dimension consisted of $TD = 256$ points with increment $\Delta t_1 = 200\ \mu s$, corresponding to a demonstration evolution time of $T_1 = 5.1$ ms.

For each RF amplitude setting and phase-cycle element, a complex free-induction decay (FID) was calculated by projecting the final density operator onto transverse $^{15}\mathrm{N}$ magnetization,

$$s(t_1) = \mathrm{Tr}(\rho N_x) + i\,\mathrm{Tr}(\rho N_y).$$

Prior to Fourier transformation the simulated FIDs were multiplied by an exponential window corresponding to line broadening $LB = 16$ Hz. The reduced-dimensionality spectrum was obtained by Fourier transforming the $t_1$ dimension and combining phase-cycle elements by subtraction.

In this framework the resulting reduced-dimensionality spectral slices correspond to averaged $^{15}\mathrm{N}$ evolution

$$\exp\left[-t_1 \frac{(1+\alpha)}{2} \omega\right],$$

with observed $^{15}\mathrm{N}$ frequency

$$\nu_N = -\frac{1+\alpha}{2}\omega, \qquad \omega = (o_N - N_{\mathrm{Hz}}),$$

where the negative sign reflects the negative gyromagnetic ratio of $^{15}\mathrm{N}$. The $^{13}\mathrm{C}$-dependent splitting along the reduced-dimensionality axis is given by

$$\Delta\nu_C = \alpha(o_{CA} - C_{\mathrm{Hz}}).$$

All simulations were performed using Python with explicit density-matrix propagation.

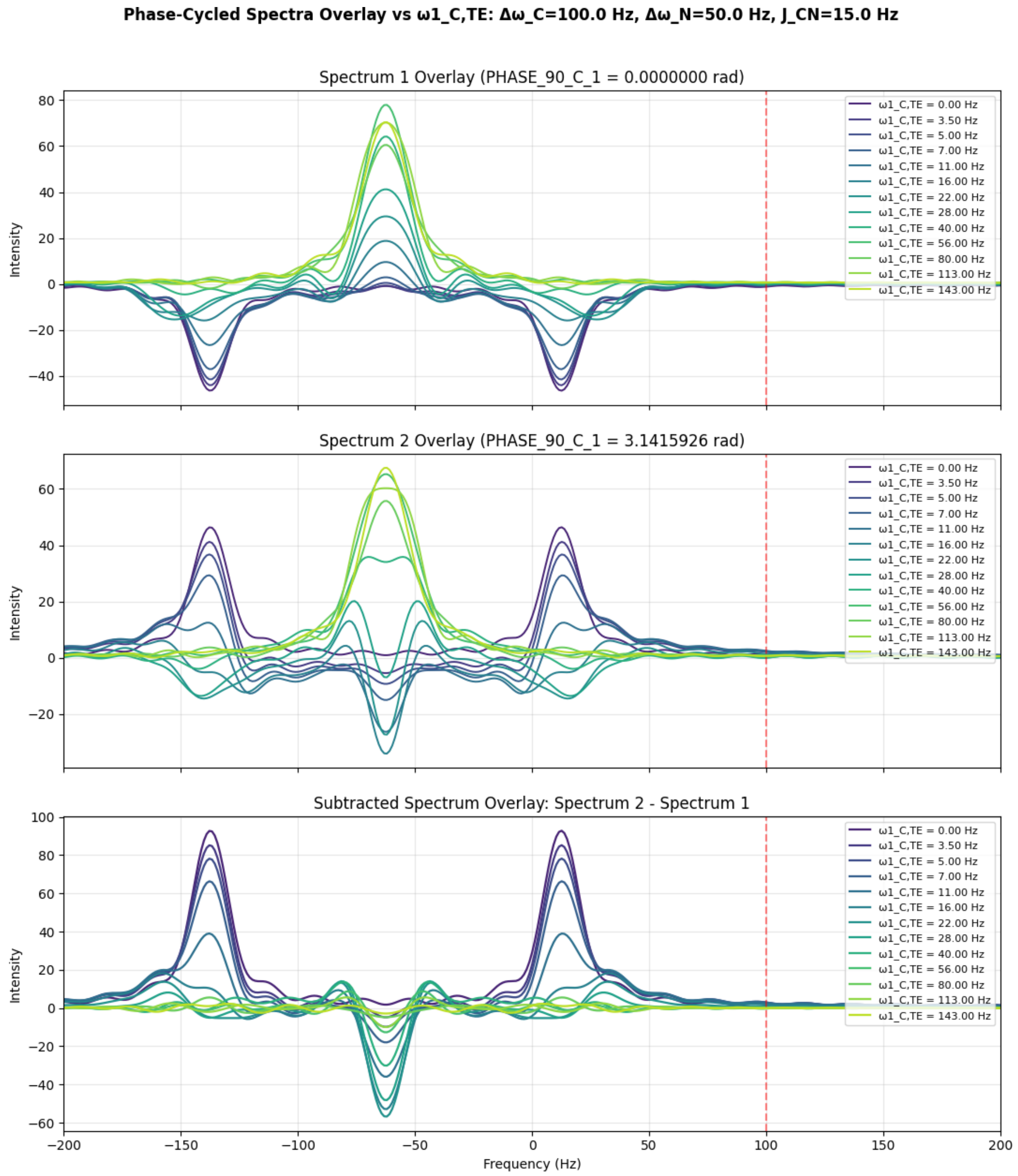


**Figure** 3S. Simulation of reduced-dimensionality (RD) one-dimensional spectra obtained from density-operator propagation of a coupled $^{13}$C, $^{15}$N two-spin system under phase-modulated weak RF irradiation. A series of spectra was calculated while varying the RF field amplitude $\omega_{1\_C}$ from 0 to 143 Hz during the transfer period and $\omega_{1\_N}$ is set to 0 Hz. Top panel: spectra obtained with the first $^{13}$C pulse phase $\phi^{(1)}_{90^{\circ}C} = 0\ rad$. Middle panel: spectra obtained with the phase-cycled element $\phi^{(1)}_{90^{\circ}C} = \pi\ rad$. Bottom panel: difference spectra obtained by subtracting the two phase-cycle elements yielding the reduced-dimensionality spectral response. The colored traces correspond to different RF amplitudes $\omega_{1\_C}$ illustrating how the spectral pattern evolves as the excitation profile of the weak RF field is shifted across the resonance. The dashed vertical line marks the reference offset of the $^{13}C$ resonance relative to the RF carrier. These simulations demonstrate the formation of RD spectral components and their dependence on RF field strength under the phase-cycled RD detection scheme.

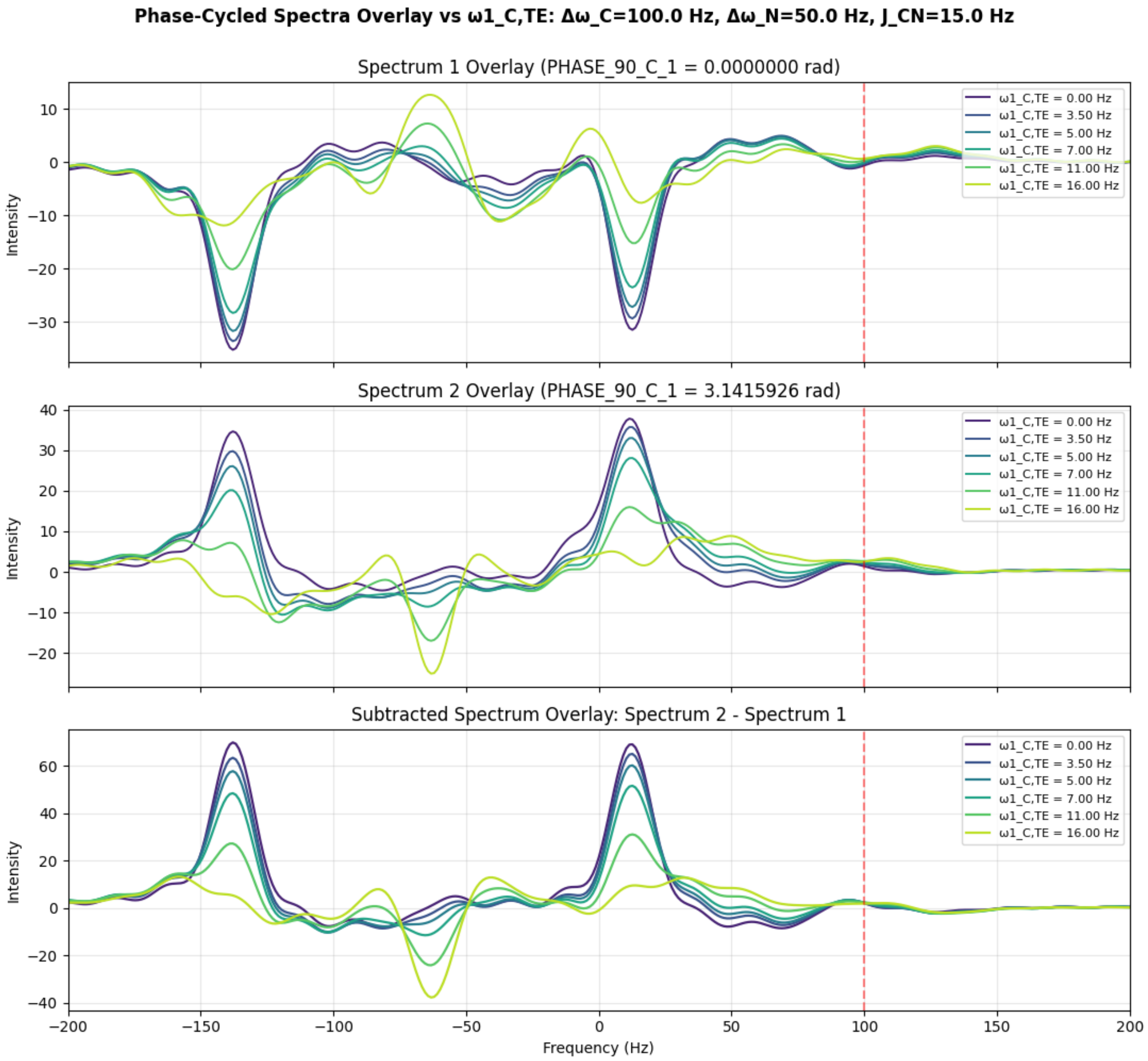


**Figure** 4S. Simulation of reduced-dimensionality (RD) one-dimensional spectra obtained from density-operator propagation of a coupled $^{13}C$, $^{15}N$ two-spin system under phase-modulated weak RF irradiation. All settings are as in Figure 3S (above) except that the series of spectra was calculated while varying the RF field amplitude $\omega_{1\_C}$ from 0 to 16 Hz during the transfer period and $\omega_{1\_N}$ is set to 5 Hz illustrating the combining effects of irradiation on both channels may significantly increase selectivity of peak amplitude attenuation.

#### 4.2. Combined weak $^{13}$C and $^{15}$N RF irradiation for highly selective peak attenuation

To investigate the selectivity of weak RF irradiation in reduced–dimensionality experiments, we simulated one–dimensional spectral responses using density–operator propagation of a coupled $^{13}$C–$^{15}$N two–spin system. The simulations followed the procedure described above, including explicit propagation of the density operator under the static Hamiltonian and time–dependent RF fields.

In addition to weak RF irradiation applied on the $^{13}$C channel during the transfer period, an additional continuous RF field was applied simultaneously on the $^{15}$N channel with extremely low RF amplitude ($\omega_{1,N} = 5$ Hz). The RF amplitude on the $^{13}$C channel was varied over a range of values, while all other parameters remained identical to those used in the reference simulations (chemical shift offsets $\Delta\omega_C = 100$ Hz and $\Delta\omega_N = 50$ Hz, scalar coupling $J_{CN} = 15$ Hz). Figure 4S demonstrates that a combination of $^{15}$N and $^{13}$C irradiations with the field strength only $\omega_{1,N} = 5$ Hz and $\omega_{1,C} = 11$ Hz might completely attenuate the corresponding RD peak.

The purpose of this simulation was to illustrate that the simultaneous application of weak RF irradiation on both nuclei dramatically enhances the spectral selectivity of peak attenuation. Two mechanisms contribute to this effect. First, the RF power applied on the $^{15}$N channel is extremely small ($\omega_{1,N} = 5$ Hz), producing a very narrow excitation profile in frequency space. Such low-power irradiation perturbs only spins whose resonance frequencies fall very close to the RF carrier, minimizing off-resonance effects on distant peaks.

Second, weak RF irradiations on the $^{13}$C and $^{15}$N channels produce attenuation profiles that are effectively orthogonal in the two frequency dimensions of the coupled spin system. The $^{13}$C irradiation modulates the magnetization transfer pathway as a function of the $^{13}$C offset, while the $^{15}$N irradiation introduces an additional selective perturbation along the $^{15}$N frequency coordinate. As a result, attenuation of the observed signal becomes strongest at the intersection of the two narrow excitation profiles defined by the $^{13}$C and $^{15}$N RF fields.

This combined irradiation therefore produces a highly localized attenuation region in the effective two–dimensional frequency space of the coupled spin system. In practice, this behavior allows selective suppression of signals from resonances located near the intersection of the two excitation profiles, while leaving other peaks largely unaffected. Such orthogonal selectivity may be exploited for targeted attenuation or selective removal of specific resonances in reduced–dimensionality experiments, particularly when minimal perturbation of neighboring peaks is required.
